# Biologically Inspired Digital Histology for Deep Phenotyping of Placental Composition Changes Across Major Lesion Types

**DOI:** 10.64898/2025.12.22.693945

**Authors:** Emma Clare Walker, Claudia Vanea, Karen Meir, Drorith Hochner-Celnikier, Hagit Hochner, Triin Laisk, Cecilia Lindgren, Craig A. Glastonbury, Linda M. Ernst, Christoffer Nellaker

## Abstract

Placenta pathology provides diagnostic insights for understanding pregnancy complications and guides maternal and perinatal care. While placental abnormalities reflect both acute and chronic maternal-fetal health states, the organ’s spatial and temporal heterogeneity poses significant challenges for systematic histological analysis. To date, there has been no analysis of cell populations and their relationship to placental lesions due to the scale of annotations required by expert pathologists. The Histology Analysis Pipeline.PY (HAPPY), previously published by our group, is a hierarchical deep learning approach for quantitative single-cell resolution analysis of H&E slides. Here we apply HAPPY to analyse 130 slides from 62 live full-term pregnancies across healthy controls and four common placental lesion types (infarction, perivillous fibrin, avascular villi, and intervillous thrombosis). We computed cell-type and tissue-structure compositions, established the expected range of healthy variation, and quantified slide-level deviation from this reference using compositional data analysis. Our results reveal significant cellular composition differences between histologically normal and lesioned placentas, including increased extra-villous trophoblasts and leukocytes, coupled with decreases in Hofbauer cells. These changes accompanied distinctive tissue microstructural alterations, particularly increased fibrin deposition and changes to the villous structures. The magnitude of compositional deviation increased with infarction size but not with intervillous thrombosis. Importantly, many differences extend beyond visibly affected areas, indicating organ-wide adaptive responses rather than purely discrete focal pathologies. This quantitative characterisation provides insights into relationships between specific pathologies and placental structure, demonstrating the potential of AI-based methods to enhance conventional histopathological assessment for research and clinical practice.

## 1. Introduction

The placenta is a complex, dynamic organ that serves as the maternal-fetal interface during pregnancy. This temporary organ facilitates essential physiological processes including nutrient and waste exchange, gas exchange, and immune regulation between mother and fetus. The placenta’s cellular and structural composition change throughout pregnancy to adapt to the needs of a growing fetus [1] and abnormalities in structure can significantly impact fetal development and pregnancy outcomes [2, 3]. Pathologies can manifest as placental lesions which are known to be associated with both maternal and fetal health outcomes, and can provide important indicators of recurrence of risk in future pregnancies [4, 5, 6]. Lesions frequently co-occur, with multiple lesions associated with worse pregnancy outcomes than single isolated lesions [7, 8, 9].

Despite the clinical importance of placental pathology for diagnosing abnormalities, significant barriers limit its use in routine clinical practice. Recent reviews have identified key obstacles including variability in describing histology findings, a shortage of perinatal pathologists, and lack of clinical understanding of placental conditions [10]. Many clinical systems simply do not have the capacity to assess all placentas which meet the clinical guidelines for referral [11]. When performed, there is high inter-observer variability in histological findings and reporting [12, 13]. The Amsterdam Workshop harmonised placental pathology criteria and established terminology for four major patterns of placental injury: maternal vascular malperfusion, fetal vascular malperfusion, acute chorioamnionitis, and villitis of unknown etiology [14, 15]. However, uncertainty persists regarding specific diagnostic criteria and the combination of histological findings to classify these [13]. Therefore, practical translation of pathology findings into actionable clinical insights remains challenging.

Recent advances in digital pathology and artificial intelligence (AI) offer promising solutions to these challenges. While AI applications in pathology have primarily focused on oncological patch-classification based approaches, placental histological assessment requires specialised models. This is largely due to the placenta’s rapidly changing structure, resilience to pathology, and limited representation in foundation models training datasets. The application of digital imaging to placental pathology has recently gained recognition [16, 17], with work including classifiers for lesion patches [18] and models for cell type quantification in placental membranes in response to chorioamnionitis [19]. Our group previously developed and published HAPPY (Histology Analysis Pipeline.PY), a biologically interpretable hierarchical approach for quantifying cell types and tissue structures at single-cell resolution across whole slide images (WSI) [20]. This automated approach enables analysis of millions of cells per slide, a scale of quantification that is infeasible for human annotators. Further, a bottom-up approach is important given limited routine placental examination, which restricts understanding and data on normal variability and underscores the need for models that capture underlying biology rather than relying on just prediction of placenta-associated pathologies.

In this study we use HAPPY to find organ-wide cellular and structural changes associated with histologically abnormal slides. We use a dataset of placenta histology slides with and without lesions, collected from three institutions with a focus on four common lesion types: infarction (ischaemic injury to villous tissue), perivillous fibrin (fibrin deposition around villi), avascular villi (loss of fetal villous capillaries), and intervillous thrombosis (blood clot formation in maternal circulation spaces). We quantify the compositional changes associated with these lesion types compared to healthy controls, and use the single-cell resolution analysis to understand the biological changes that are driving differences in histologically abnormal slides. Our approach provides quantitative metrics for placental differences and reveals changes not apparent through conventional histological examination.

## 2. Methods

### 2.1. Dataset

We conducted a retrospective analysis of placental histology slides from three institutions. The dataset used in this work was obtained from placental Hematoxylin and Eosin (H&E) slides collected as part of routine clinical pathology investigation. We obtain 740 slides from 230 patients and associated filtered pathology notes, collected between 2016 and 2017 from Hadassah Medical Center, Israel (HMC); 159 slides from 41 patients, and associated free text pathology reports, collected between 2016 and 2020 from the University of Tartu, Estonia (UoT); and 22 slides from 22 patients, derived from term pregnancies with histologically normal placentas, after full gross and histological examination, collected between 2021 and 2022 from Endeavor Health, United States (EH). Our final dataset of full term parenchyma slides, with the lesion(s) of interest, consists of 130 slides from 62 patients (UoT: 8 slides from 3 placentas, HMC: 103 slides from 40 placentas, EH: 19 slides from 19 placentas). This study uses slides that were obtained from singleton pregnancies at *≥* 37 weeks gestation.

Local ethics committees at each institution approved the use of all pseudoanonymised samples and waived consent requirements (UoT approval 289/T-5 by the Research Ethics Committee of the University of Tartu; HMC approval 0735-18-HMO by the Helsinki Committee at the Hadassah Medical Center; EH exemption EH23-303 by the institutional review board at Endeavor Health.

### 2.2. Histological Preparation and Digital Imaging

Histology slides were prepared using a standard formalin fixing, paraffin-embedding, and H&E staining procedure. As per clinical guidelines, full-thickness samples including both chorionic and basal plates were sampled and 5*µm* thickness slices were generated. Slides were scanned and digitised using a Hamamatsu XR, a 3D HISTECH PANNORAMIC 250 Flash III or Aperio GT 450 scanners at x40 magnification.

### 2.3. Pathologist Annotations

Following digital image acquisition, we included parenchyma slides in this study that were either histologically normal or contained at least one lesion of interest from the following: infarction, perivillous fibrin deposition, avascular villi, or intervillous thrombosis. We excluded non-singleton pregnancies and slides with significant scanning artefacts..

Where available, pathology reports provided assessment for the whole placenta examination. Slide-specific annotations were performed by LE and CV in conjunction with pathology reports to identify which slides contained lesions and which were histopathologically normal. Estimation of lesion coverage across slides was performed by LE and ECW using binned percentage estimates.

### 2.4. Model Overview

We applied the previously published Histology Analysis Pipeline.PY (HAPPY) using pretrained models for placental parenchyma analysis from [20], where the graph structure classification model has been updated from the original paper as described in Campbell et al. [21]. HAPPY is a three stage image analysis deep learning pipeline for analysing histology images, built with a biological hierachy: 1) an object detection model for nuclei localisation, 2) an image classification model for cell classification, 3) a graph neural network (GNN) for tissue classification, model overview in Fig 1a. This allows us to construct a spatially informed cell graph across the H&E whole slide image (WSI). All inference was performed on a single NVIDIA A100 GPU.

**Figure 1:**
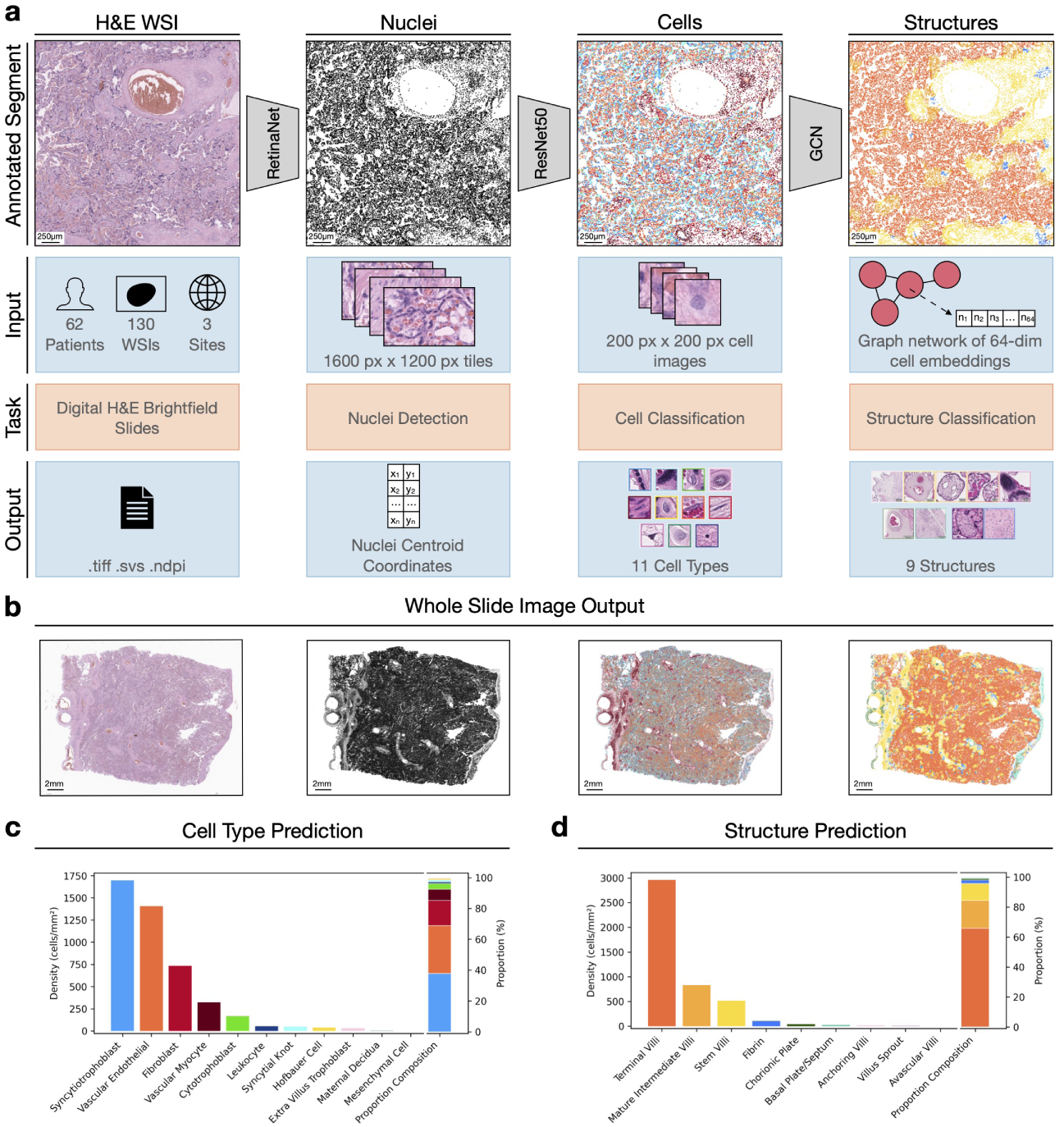
Overview of the HAPPY (Histology Analysis Pipeline.PY) workflow for placental histology analysis. (a) The three-stage deep learning pipeline processes digitalised H&E-stained whole slide images (WSI) through: nuclei detection, cell classification, and tissue structure classification. (b) Example outputs from analysis on a representative healthy term placental WSI, showing the input H&E histology image, nuclei detection, cell classification, and structure classification. (c) Predicted cell types and (d) predicted structure types, quantifying both density (cells/mm²) and percentage composition of the slide (%). WSI, whole slide image

#### 2.4.1. Nuclei Detection

The WSI is divided into patches of 1600×1200-pixels (177.44×133.08 *µm*) with a 200-pixel (22.18 *µm*) overlap for input to an object detection model (RetinaNet with a ResNet-101 backbone) to identify all nuclei within the patch. The nuclei coordinates are recorded with duplicate predictions within a 4-pixel radii removed.

#### 2.4.2. Cell Classification

The cell classification stage uses the nuclei coordinates to extract and crop a 200×200-pixel (22.18×22.18 *µm*) image centred on each nucleus. A ResNet-50 model classified cells into 11 types: four trophoblast variants (syncytiotrophoblast, cytotrophoblast, syncytial knot, extravillous trophoblast), five mesenchymal-derived cells (fibroblast, Hofbauer cell, vascular endothelial cell, vascular myocyte, undifferentiated mesenchymal cell), and two non-villous cells (maternal decidual cell, leukocyte). Post-processing isolated syncytial knot predictions are converted to syncytiotrophoblasts and group clusters to syncytial knots appropriately.

#### 2.4.3. Structure Classification

The coordinates and cell representations are used to construct a spatial cell graph. Graph nodes are the 64-dimensional embedding from the last ResNet-50 latent layer, and edges represent probable cellular interactions within tissue structures through intersection of k-nearest neighbours (k=5) and Delaunay triangulation algorithms. A ClusterGCN graph neural network with 16 GraphSAGEConv layers classifies each cell into one of nine tissue structures: five chorionic villous types (stem, anchoring, mature intermediate, terminal villi, villous sprouts), two maternal/fetal surfaces (chorionic plate, basal plate/septum), and two pathological indicators (fibrin, avascular villi).

#### 2.4.4. Data Processing

From HAPPY inference of the WSIs, we extract cell coordinates, and cell-type and structure predictions for each slide. We estimated tissue area by dividing slides into non-overlapping patches and aggregating the area of patches containing at least one nucleus prediction, excluding background regions. We calculated cell and structure distributions as both proportions (percentage composition) and densities (cells/mm²), normalising for pixel dimensions (0.1109 micrometres per pixel) and tissue area measurements. Quality control steps can be found in Supplementary Information 1.

#### 2.4.5. Quality Control

We implemented multiple quality control measures throughout our workflow. First, we removed slides with observationally poor scanning quality, including those with scanning artefacts (such as duplicated regions) identified through manual inspection. Second, we excluded all cells classified as chorionic plate or basal plate from analysis to focus on parenchymal regions and control for interinstitutional sampling variations. Third, following Aitchison distance testing (described below), we removed outlier slides showing high distributional differences compared to healthy controls that, upon manual inspection, contained scanning artefacts or processing errors. These quality control steps resulted in the exclusion of five slides from the final dataset.

### 2.5. Statistical Analysis

Statistical significance was assessed using multiple statistical methods: Kolmogorov-Smirnov tests were used to evaluate differences in Aitchison distance distributions between groups of slides (described in Section 2.5.1), Mann-Whitney U tests were applied to individual cell-type and tissue-structure density measurements. Pearson correlation coefficents were calculated to assess the relationship between lesion coverage and Aitchison distance, as well as the correlation between the average Aitchison distance of slides with and without lesions present from the same patient. Local autocorrelation and heatmaps were used the for visualisation of cellular and structure spatial distributions (described in Section 2.5.2). Bonferroni correction was applied where appropriate to adjust for multiple testing.

#### 2.5.1. Aitchison Distance for Quantifying Compositional Differences

Aitchison’s distance, which provides a distance metric for measuring the difference between two compositional vectors, is used to quantify how much each slides’ cell-type or structural composition deviated from a healthy reference composition. For a compositional vector **p** = (*p*_1_*, . . ., p_k_*), , where *p_i_* denotes the proportion of the *i*-th component and Σ*^k^_i=1_ p_i_* = 1, we first applied the centred log-ratio (CLR) transform:

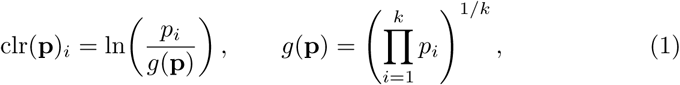

where *g*(**p**) is the geometric mean of the *k* compositional parts. The Aitchison distance (*d_A_*) between two compositions **p** and **q** is then calculated as the Euclidean distance between the CLR-transformed vectors:

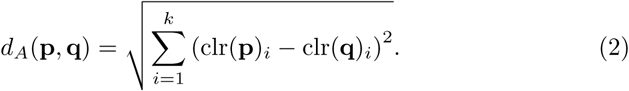

To characterise normal spatial heterogeneity, each healthy control slide (n=19) was segmented into three regions along an axis from the chorionic plate to basal plate, and percentage compositions of cell and structure types were computed for each region. We performed bootstrap resampling of the healthy training regions (n=10,000), computing the *d_A_* between randomly sampled region pairs to obtain a measure of the expected distribution of healthy compositional variation (sampling strategy, Supplementary Fig. 2). The mean healthy reference composition was calculated from the control slide compositions. Significance thresholds were defined as the 95th percentile of the healthy bootstrapped *d_A_* values (cell-type threshold *d_A_* = 3.32; structure-type threshold *d_A_* = 3.90).

For each test slide, we computed the *d_A_* between its cell-type or structure composition and the healthy reference composition. Slides with distances exceeding the corresponding significance threshold were considered to have significant compositional deviation. Error bars on the percentage of significance slides are determined by leave-one-out resampling where the average is recalculated after removing one slide in each iteration to determine the stability of the average (error bars reflect the variability across these estimates).

#### 2.5.2. Spatial Analysis with Local Autocorrelation

A local autocorrelation metric is used to quantify and visualise the distribution of cell types and structures across slides. We divided slides into 150×150 grid squares, calculating local autocorrelation at each grid point *i* as:

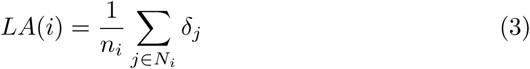

where *LA*(*i*) represents the local autocorrelation value at grid point *i*, *n_i_* represents the number of cells within a 200*µm* radius from grid point *i*, *N_i_* denotes the set of all cells within this 200*µm* radius, and *δ_j_* serves as a binary indicator (1 if cell *j* belongs to the target type, 0 otherwise). Grid squares containing fewer than 5 cells were excluded from analysis and appear as transparent in visualisations. We applied Gaussian interpolation smoothing for visualising the heatmap.

## 3. Results

### 3.1. Quantification of cellular and structural composition in placental histology

Our dataset is comprised of 130 full-term placental parenchyma WSI slides from 60 patients across three institutions. This contains both placenta WSIs from healthy uncomplicated pregnancies (control) slides (n=19), slides with lesions present (n=95), and slides that appear histopathologically normal from placentas with lesions present elsewhere (n=16), Supplementary Fig. 1 summarises the selection criteria. All pregnancies resulted in singleton, live births. We use HAPPY to extract quantitative cellular and tissue structure metrics from our dataset. The pipeline, shown in Fig 1a, is used to process WSIs through a three stage biologically-inspired hierarchical approach of nuclei detection, cell classification and graph-based tissue structure classification. Across our slides, we identified 165,327,306 nuclei (1, 132, 379 *±* 425, 906 mean nuclei per slide), classified cells into 11 cell types, and tissue structures into 9 categories covering the relevant parenchyma cell types and structures. After applying HAPPY, each nucleus has an associated cell-type and structure prediction from which we calculate density measurements and relevant compositions.

We show an example output from a healthy slide (H&E), visualising the corresponding detected nuclei, classified cells, and tissue structures in Fig 1b. Cell type and structural composition align with biological expectations for a healthy term placenta, Fig 1c. We observe syncytiotrophoblasts are the dominant cell type with relatively few cytotrophoblasts, and terminal villi comprise the highest density of the tissue structures, with negligible avascular villi.

### 3.2. Compositional Differences of Slides with Lesions Present

We first examined slides containing four major placental lesion types (Fig 2a): infarction (n=34), perivillous fibrin (n=45), intervillous thrombosis (IVT) (n=37), and avascular villi (n=15), comparing these against histopathologically normal control slides (n=19) (Supplementary Table 1) focusing on the parenchyma region to reduce sampling bias (Fig 2b). Slides can have multiple lesions (single lesion n=64, multiple lesions n=31) and this is defined as a separate category for the quantitative analysis.

**Figure 2:**
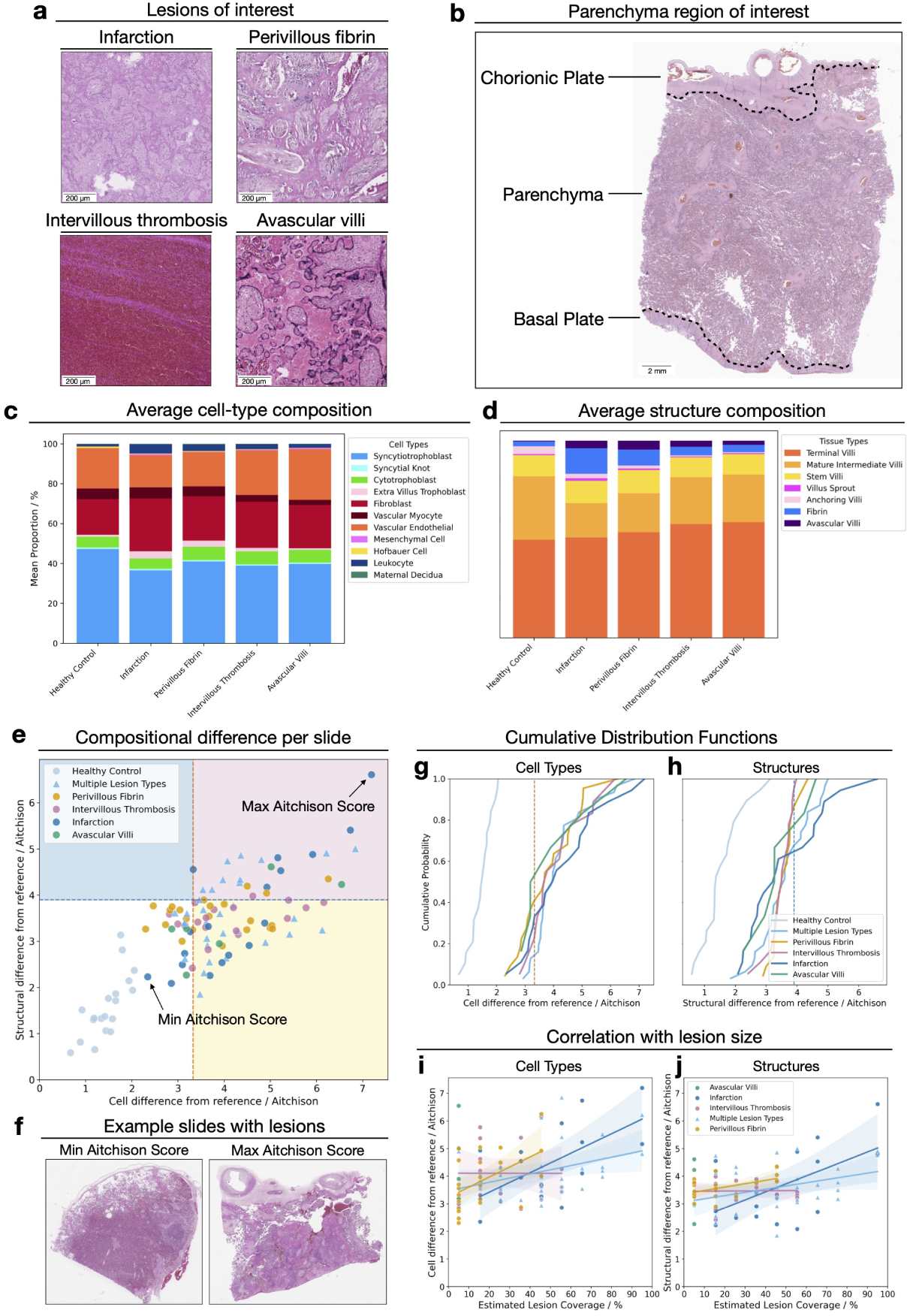
(a) Example histological segments of the four lesions of interest: infarction, perivillous fibrin deposition, intervillous thrombosis and avascular villi. (b) H&E image of a healthy term placenta showing the chorionic plate, parenchyma and basal plate. (c-d) Predicted cell-type (c) and structural (d) compositions averaged from the parenchyma region in healthy slides and slides with lesions. (e) Distribution of Aitchison distances per slide, quantifying deviation from the healthy reference composition. Slides are coloured by lesion category and regions with significant cellular (yellow), structural (blue), or both (pink) compositions differences are indicated. (f) H&E images of lesioned slides with the minimum (left image; small localised infarction) and maximum (right image; slide is entirely infarcted) Aitchison distances. (g-h) Cumulative distribution functions of the Aitchison distance for each lesion groups cell-type (g) and structural (h) composition, all significantly different than the healthy control (Kolmogorov-Smirnov test, *p <* 0.001). (i-j) Correlation between estimated lesion coverage and Aitchison distance for cellular (i) and structural (j) correlation, with trend lines and 95% confidence intervals.

When we consider the aggregated average cell-type and structure composition for slides containing any of the four lesion types, there are differences in the overall composition of healthy slides against slides with lesions (Fig 2c and Fig 2d). Despite typically comprising a small region of the slide, these shifts are still detectable at the average whole slide-level composition compared to controls.

We define each slide’s compositional difference from the healthy reference composition using Aitchison’s distance between two slides (see Methods 2.5.1). This quantifies the fold change in each component of the cell-type or structure composition in comparison to the healthy composition. This approach identified extreme outliers, which on further manual inspection we determined to have technical artifacts and 5 slides were removed from further analysis.

All control slides fall within the bootstrapped significance thresholds, yet the intra-slide compositional differences observed is greater than those observed across whole-slide tests (Supplementary Fig. 2), reflecting both spatial heterogeneity and the constraints of a small healthy reference set. Despite the variation within healthy tissue, we detect slides with lesions that fall outside expected healthy range, with cell-type and structure composition differences shown in Fig 2e. We find that slides with an infarction present have the widest range of compositional differences (38.9% of infarction slides exceed both cellular and structural significance thresholds, 72.2% of slides have a significant difference in the cell-type composition, 38.9% have a significantly different structural composition). In contrast, slides with IVT showed the most constrained compositional changes to the wider structures present on the slide (5.56% of slides exceed both thresholds, 77.8% of cell-type compositions, 5.56% of structural compositions). We present the details results across all lesion categories in Supplementary Fig. 3.

To understand whether groups of slides with lesions have significant composition differences compared to the healthy slides, we compare the cumulative distribution functions (CDF) of the Aitchison distance values of lesioned slide group with the healthy control group, testing with Kolmogorov-Smirnov (KS) tests (Fig 2g-h). All groups of lesion types have significantly different cell-type composition profiles compared to healthy controls (*p <* 0.001). Analysing the structural composition, we found slides with infarction (*p <* 0.001) and perivillous fibrin (*p <* 0.01) showed significant deviations, whilst the distribution of Aitchison distance values for slides with avascular villi and IVT did not differ significantly from the healthy control group.

To test if batch effects were driving observed changes we did a sub-analysis of slides with an infarction to understand if the Aitchison metric varies between sites. As the values from both sites cluster together we conclude that a batch effect is not driving the changes (Supplementary Fig. 10).

Next we performed a manual estimation of the proportion of a slide covered by lesions and compared this to the Aitchison distance to see if the size of a lesion is correlated with differences in the cell or structural composition of a slide (Fig 2i,j). For slides with an infarction present, we find a significant correlation between the size of the lesion and the Aitchison distance (Spearman rank coefficient, infarction: r=0.61 *p <* 0.05). We do not detect a significant change in the structural composition (r=0.42 *p* = 0.41). Slides with perivillous fibrin, intervillous thrombosis, or multiple lesions types did not have significant correlation between the cellular or structural composition deviation and the estimated area of the slide covered by the lesions (perivillous fibrin, cellular: r=0.48 *p* = 0.11, structural: r=0.33, *p* = 0.64, multiple lesion types, cellular: r=0.34, *p* = 0.33, structural: r=0.32, *p* = 0.40, IVT, cellular: r=-0.016, *p* = 1.0, structural: r=-0.078, *p* = 1.0). We do not measure the correlation for slides with avascular villi as there is only *<* 10% coverage of any slide with avascular villi. An examination of individual slides further supports this pattern (Figure 2f). We show the slides with a lesion present with the maximum and minimum Aitchison score. Both slides have an infarction present, however the slide with the largest cell-type and structural composition difference is predominately covered by an infarction. Conversely, the slide with the minimum combined score has a small localised infarction present.

### 3.3. Deep Phenotyping Indicates Compositional Differences in Slides where the Lesion is Not Apparent

To investigate whether focal lesions induce compositional changes that extend throughout the placenta beyond visually affected regions, we analyse slides where both histologically normal (‘no apparent lesion’) and lesion-containing slides were available from the same patient (n=13 patients) (Fig 3a). We detect a change in the composition of the cell-types and structures even in those slides which have been determined to not have an apparent lesion present by a pathologist (Fig 3b,c). Among slides with ‘no apparent lesion’, 60% have significantly different cellular compositions from healthy controls slides, and 13.3% have significantly different structural compositions (Fig 3d). We present the detailed results in Supplementary Fig. 4.

**Figure 3:**
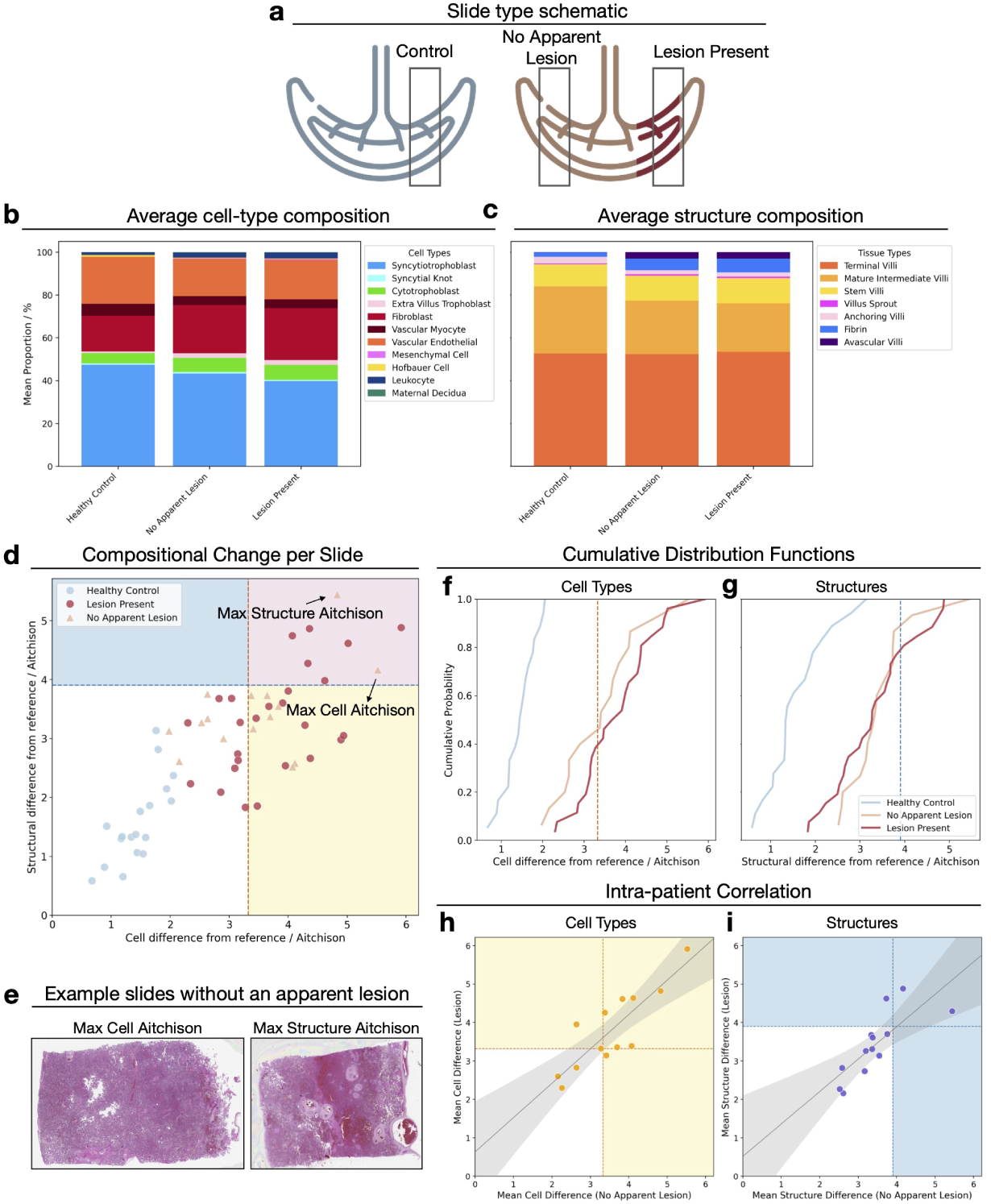
(a) Schematic of sampling regions from healthy control placentas and placentas with lesions, showing slides with no apparent lesion and those with the lesion present. (b-c) Predicted cell-type (b) and structural (c) compositions averaged across these groups. (d) Distribution of Aitchison distances per slide, quantifying deviation from the healthy reference composition. Slides are coloured by slide type, and regions with significant cellular (yellow), structural (blue), or both (pink) compositions differences are indicated. (e) H&E images of slides with no apparent lesion showing the highest Aitchison distance for cell composition (left; histologically normal in appearance) and the highest distance for structural composition (right; abnormal-appearing slide with numerous stem villi but no discrete lesion). (f-g) Cumulative distribution functions of Aitchison distances for each lesion group for cell-type (f) and structural (g) composition, all significantly different than the healthy control (Kolmogorov-Smirnov (KS) test, *p <* 0.001). (h–i) Intra-placental correlations: each point represents a patient and shows the relationship between their average Aitchison distance on no-apparent-lesion slides and their slides with lesions, for cellular (h) and structural (i) composition, with trend lines and 95% confidence intervals.

We consider the CDFs of the Aitchsion distances for healthy controls, slides with ‘no apparent lesion’, and those with lesions to assess how the presence of a lesion affects compositional distributions (Fig 3f,g). We find that the CDF of both the cell-type composition and the structure composition for slides ‘no apparent lesion’ are significantly different from the healthy control group (KS test, *p <* 0.001 for all tests). Examining the histopathologically normal slides with extreme deviations in Fig 3e, there is no identifiable abnormalities to a casual or expert human interpreter for the slide with the maximum cell composition difference. Conversely, for the structural example, it is apparent to a human observer that the slide is abnormal and numerous stem villi present but there is not a classifiable lesion present.

Next, we sought to determine whether there was a correlation between the compositions from slides within the same placenta to understand if the lesion affects distal slides within the same placenta (Fig 3h, i). We find that slides with lesions and slides without lesions from the same placenta have significantly correlated average Aitchison distances based on the cell-type compositions (Spearman rank coefficient, r = 0.84, *p <* 0.001). Similarly, we find significant correlation in the structure compositions within individuals placentas (r = 0.87, *p <* 0.001). These findings could be interpreted as evidence for systemic responses throughout the placenta when lesions are present. Alternatively, this correlation could reflect inherent patient-specific compositional characteristics that predispose certain placentas to lesion development.

### 3.4. Specific Cell and Structure Density Changes Driving Composition Changes

To understand what is driving the compositional differences, we identify several key cell populations that show consistent patterns in response to injury and find these changes match our current understanding of placental biology (Fig 4). Leukocyte density is significantly increased across all lesion categories, including in slides where the lesion is not present (*p <* 0.01) (Fig 4a). Hofbauer cells demonstrate the opposite pattern, with significant depletion across all lesion types and significant reduction in ‘no apparent lesion’ slides compared to healthy controls (*p <* 0.001) (Fig 4b). Extravillous trophoblast (EVT) density is significantly increased across all lesion types compared to healthy controls. EVT density was significantly elevated in ‘no apparent lesion’ slides compared to controls (*p <* 0.01), and further increased in ‘lesion present’ slides compared to both controls (*p <* 0.001) and ’no apparent lesion’ slides (*p <* 0.01) (Fig 4c). We present full cell analyses across lesions and slide types in Supplementary Fig. 5,6. Magnitudes shown in supplementary table

**Figure 4:**
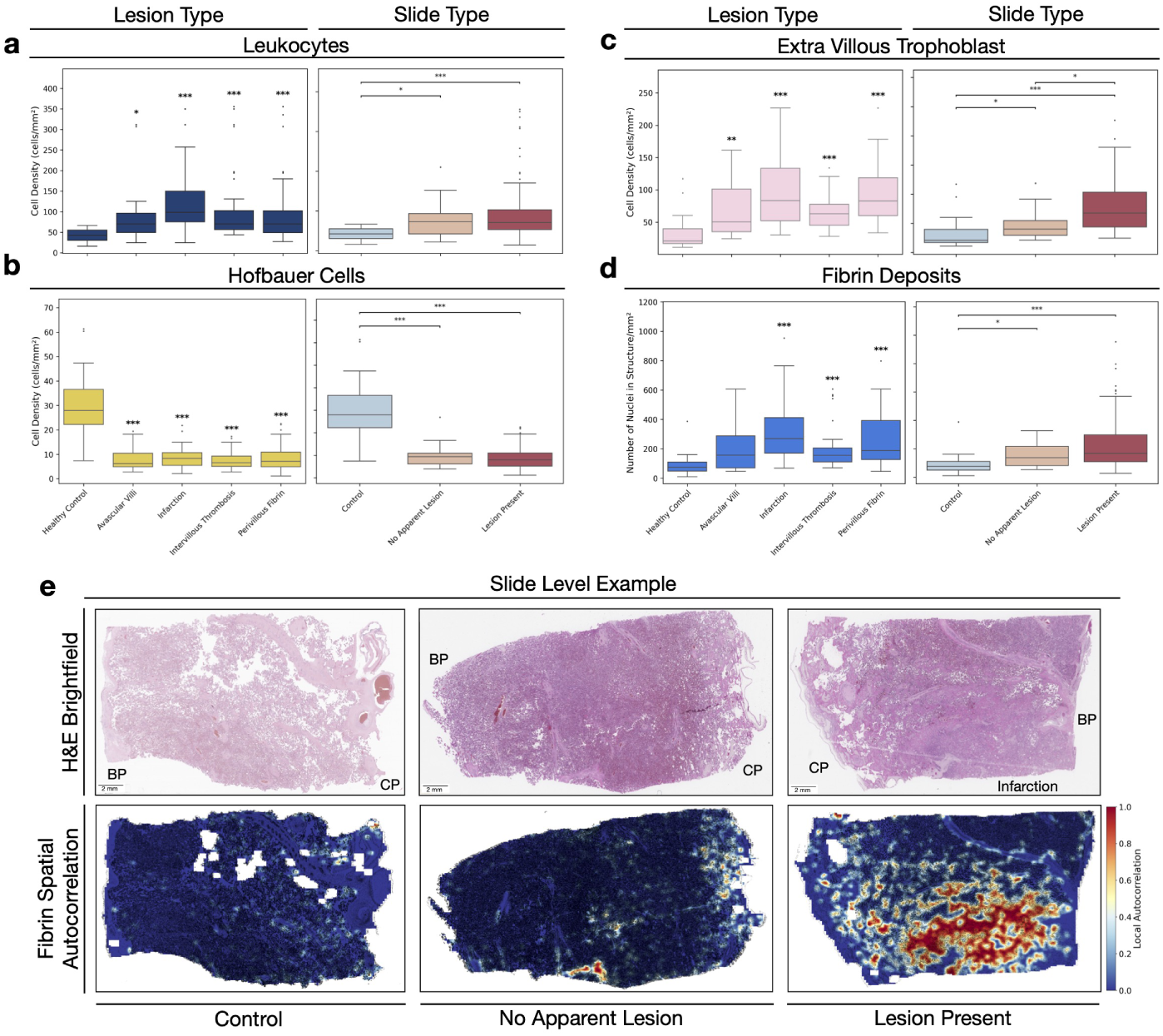
Specific cellular and structural density changes underlying placental composition changes. Cell densities (cells/mm^2^) across lesion types (left) and slides types (right) for (a) leucocytes, (b) Hofbauer cells, (c) extravillus trophoblasts, and (d) fibrin deposits. Statistical comparisons performed using Mann-Whitney U test with Bonferroni correction, with lesion categories tested against healthy control group and slide types tested pairwise. ∗ ∗ ∗*p <* 0.001, ∗ ∗ *p <* 0.01, ∗*p <* 0.05. (e) Representative H&E slide examples (top) with spatial distribution of fibrin, measured by local autocorrelation (bottom). Red regions indicate clustered fibrin deposits, whereas blue regions reflect low or dispersed fibrin. Control slides and slides without an apparent lesion show typical distribution, whilst lesion-present slides demonstrate increased fibrin deposition concentrated around the infarction site. BP: Basal plate; CP: Chorionic plate.

At the structural level, we observe significant increases in the density of fibrin deposits across all lesion types, except in slides with avascular villi, compared to healthy controls (Fig 4d). Fibrin density is also significantly increased in slides with no apparent lesions compared to healthy controls (*p <* 0.01). We present full tissue structure analyses across lesions and slide types in Supplementary Fig. 7,8.

Finally, we use spatial local autocorrelation analysis to examine the distribution of cell and structure types across slides, with a particular focus on fibrin deposits (Fig 4e). We demonstrate extensive fibrin clusters concentrated around infarction areas, suggesting fibrin distribution could guide lesion identification. In contrast, when examining fibrin localisation in ‘no apparent lesion’ and the healthy controls slides we see a more dispersed pattern. All WSIs with infarction and paired ‘no apparent lesion’ slides are visualised in Supplementary Fig. 9.

## 4. Discussion

In this study, we used a hierarchical modelling approach to identify biological changes in placenta histopathology slides in response to injury, applying the HAPPY pipeline to automatically quantify cellular and structural composition in a cross-institutional dataset. We demonstrate the utility of the approach on slides obtained through standard clinical pathology workflows, showing that slides from abnormal placentas are associated with quantifiable shifts in both cell-type and structural composition across WSIs of the parenchyma. In placentas with lesions, we detect compositional differences in slides even where the lesions are not histologically apparent to a pathologist.

We define a distance metric between two compositions to quantify the magnitude difference in a slide’s composition. In healthy placenta slides, intra-slide differences in structural composition are larger than the differences observed between whole-slide compositions across the healthy reference set, reflecting spatial heterogeneity within individual placentas. When considering cellular and structural composition across WSIs, we detect significant differences in slides with lesions compared to healthy controls, matching known histological changes [22]. Notably, more slides show significant cellular than structural deviations, which may reflect the inherent heterogeneity of villous tree sampling and limited power to overcome this heterogeneity. As all our samples resulted in live births, our slides may have a limited representation of tissue structure remodelling associated with more severe injury. Taken together, the results suggest a cellular response that may precede or mitigate structural remodelling, but longitudinal data are required to confirm causality. Future work should investigate a full spectrum of outcomes to study if limited changes to the structural composition differences here reflect compensation mechanisms that maintain placental function even in the presence of lesions [23].

For infarction, we find that the size of the lesion correlates with the compositional difference, conversely we do not see this correlation for slides with an IVT. This aligns with reported histological findings: infarction involves widespread ischaemic damage requiring extensive compensatory responses, whilst thrombotic lesions typically maintain well-defined borders with minimal surrounding tissue involvement [24, 25, 26, 27].

We also find we have power to detect differences in slides where the lesion is not apparent to a pathologist. As HAPPY quantifies the composition of slides rather than classifying specific lesion types directly, it allows for detection of subtle changes in the parenchyma, separate from primary lesion features. We demonstrate that placental lesions have detectable effects distally from the lesion site: compositions showed significant correlations between lesion and histologically normal regions within the same placentas. This may suggest progressive adaptation, such as an increase in extra-villous trophoblasts, rather than the placenta being a static organ with excess capacity [28] [23]. Such adaptive processes may help explain why as many as half of ”healthy” pregnancies appear to have histological abnormalities in the placenta [29] [30]. Though this may also suggest inherent correlation between the structures and cellular composition of placentas throughout the organ.

Cellular changes correspond to established placental pathophysiology [31] [32] [22] [33]. Systematic increases in extravillous trophoblasts across all lesion types reflect the compensatory vascular remodelling responses documented in hypoxic injury patterns [34] [35]. Progressive leukocyte infiltration aligns with expectations of an organ-wide inflammatory response to tissue injury [36] [37]. Similarly, excessive fibrin deposition aligns with expected pathological findings of patterns of injury [38], with studies indicating structural support mechanisms associated with immune processes [39] [40]. In addition, decreases in Hofbauer cell densities across lesion types align with previous reports [41] [42] [33] where prior studies have proposed that such reductions could reflect immune-mediated depletion or migration to fetal structures for local inflammatory responses [41]. While this study demonstrates the biological validity and potential clinical utility of the HAPPY pipeline, it is limited by a few key challenges. Most importantly, the sample sizes, particularly for healthy controls and paired patient analysis are small. We restricted our dataset to full-term samples to control for gestational-age–related compositional differences. However, all included cases were from live births and may not capture the full range of histological features. Most samples (UoT + HMC) were obtained from routine pathology submissions, which further restricted the availability of full-term healthy controls. Histological sampling of placental tissue is also spatially inconsistent between individuals meaning routinely collected slides may not be representative of the whole underlying organ, and we do not have data on how sampling was performed. Future work should focus on obtaining more representative and detailed placental data to increase the robustness of data for downstream analysis. Finally, we show that HAPPY would be a good basis for investigating association of placental phenotypes with future maternal and neonatal health outcomes [43, 44, 45, 5]. In summary, the bottom-up approach of our pipeline allows us to identify cell-type and tissue structure changes that would be missed using direct AI lesion image classification methods and we are able to detect changes in slides typically labelled as histologically normal. Our approach is explainable, allowing for specific characterisation of the biology of the parenchyma, augmenting the work of histopathologists and supporting trust in model outputs. The use of a trustworthy, reproducible, and quantitative workflow could help address the well-described obstacles to incorporating routine placental histological analysis into clinical practice [10] [12].

## Data and Code Availability

The HAPPY codebase, training data, and trained models for placenta histology are available at (https://github.com/Nellaker-group/happy). Cell and structure proportions and densities for each WSI per lesion category is available in Supplementary Data.

Following institution policies, all requests for data curated in-house will be evaluated on a case-by-case basis to determine whether the data requested is compliant with data sharing obligations. Data can only be shared for academic research purposes and will require a material transfer agreement.

## CRediT Authorship Contribution Statement

**E.C. Walker**: Conceptualisation, Methodology, Software, Validation, Formal Analysis, Investigation, Data Curation, Writing - original draft, Visualisation. **C. Vanea**: Methodology, Software, Data Curation **K. Meir**: Resources, Data Curation. **D. Hochner-Celnikier**: Resources, Data Curation. **T. Laisk**: Resources, Data Curation. **H. Hochner**: Resources, Data Curation. **C. Lindgren**: Resources, Funding acquisition. **C. Glastonbury**: Writing - review and editing, Supervision. **L. Ernst**: Resources, Data Curation, Writing - review and editing, Supervision. **C. Nellaker**: Conceptualisation, Resources, Writing - review and editing, Supervision, Funding acquisition.

## Generative AI

During the preparation of this work the author(s) used ChatGPT Edu in order for proof reading. After using this tool/service, the author(s) reviewed and edited the content as needed and take(s) full responsibility for the content of the published article.

