## Supplementary Information for "Biologically Inspired Digital Histology for Deep Phenotyping of Placental Composition Changes Across Major Lesion Types"

Supplementary Section 1

Supplementary Section 1 – Further processing details

**1. Quality Control**

We implemented multiple quality control measures throughout our workflow. First, we removed slides with observationally poor scanning quality, including those with scanning artefacts (such as duplicated regions) identified through manual inspection. Second, we excluded all cells classified as chorionic plate or basal plate from analysis to focus on parenchymal regions and control for inter-institutional sampling variations. Third, following Aitchison distance testing (described below), we removed outlier slides showing high distributional differences compared to healthy controls that, upon manual inspection, contained scanning artefacts or processing errors. These quality control steps resulted in the exclusion of five slides from the final dataset.

**Supplementary Information**

Supplementary Figures 1-10

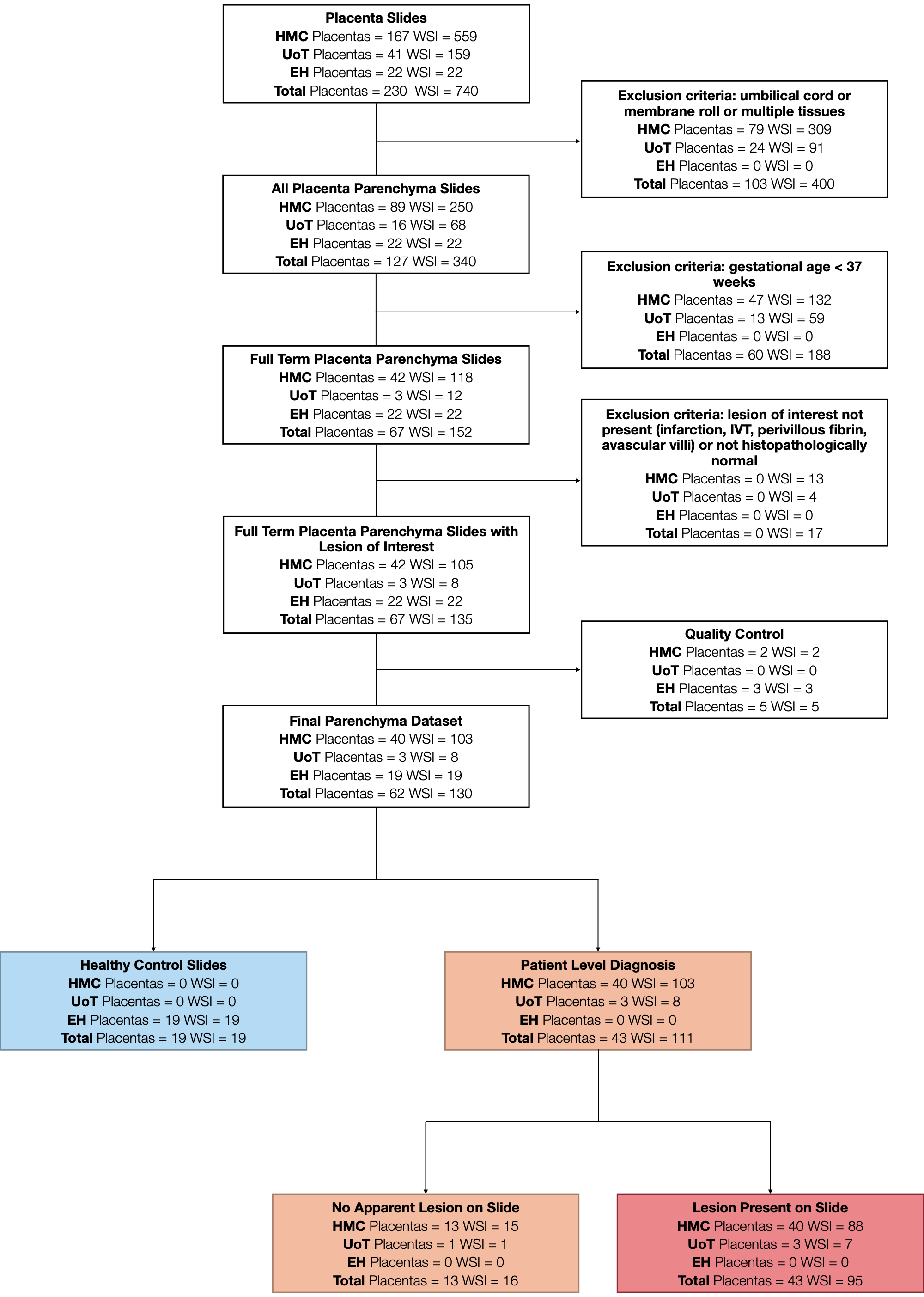

**Supplementary Figure 1:** Flow chart of exclusion criteria to select slides for final analysis. HMC = Haddash Medical Centre, UoT = University of Tartu, NUH = Northshore University HealthSystem

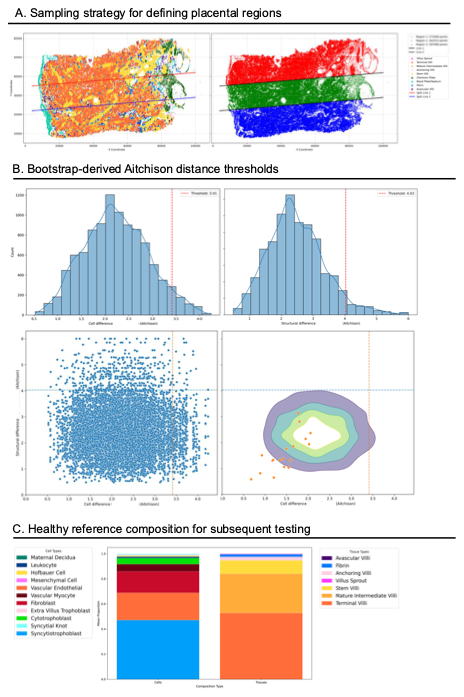

**Supplementary Figure 2:** (A) Example sampling strategy used to define anatomical regions within placental whole-slide images. Segmentation outputs and region boundaries are shown for one representative slide. (B) Bootstrap testing of regional compositions to characterise the variation in Aitchison distances among healthy slides. Histograms show the distribution of bootstrap-derived distances (n = 10,000) for cell-type (left) and structural (right) compositions, with red lines indicating the 95th-percentile significance thresholds. Scatter and contour plots depict the joint distribution of cellular and structural distances from pairwise bootstrap samples. Points overlaid on the density plot represent individual healthy slides, illustrating that their Aitchison distances to the healthy reference are smaller than the intra-slide variation observed in the bootstrap samples. (C) Reference compositions used to calculate Aitchison distances per slide, computed as the mean cell-type (left) and structural (right) compositions across healthy control slides. These reference values serve as the baseline for assessing composition differences across lesion categories and slide types.

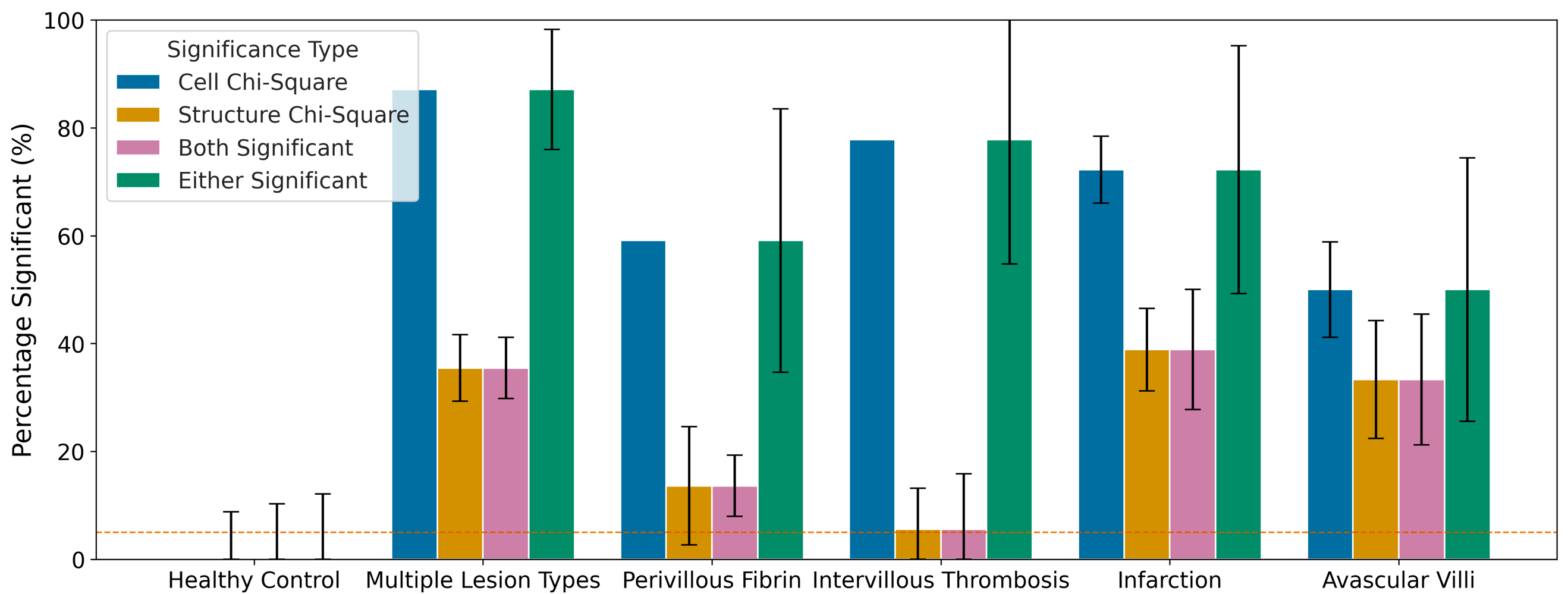

**Supplementary Figure 3:** Percentage of slides within each lesion category that show significant composition differences relative to the healthy reference. Bars indicate the proportion of slides exceeding the Aitchison-distance significance thresholds for cell-type, structural, both, or one slide having significance with either composition. Error bars represent the uncertainty estimated by recalculating the percentages leaving out on slide at a time, showing how sensitive the results are to the inclusion of any individual slide.

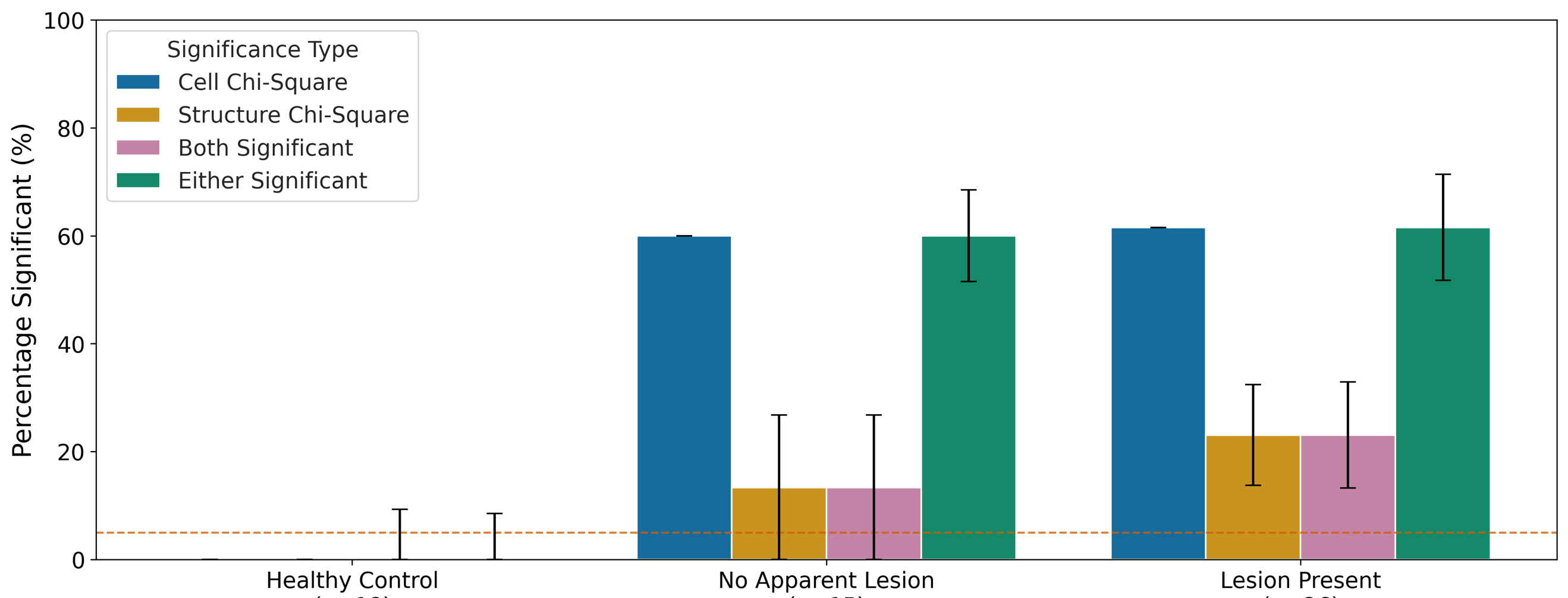

**Supplementary Figure 4:** Percentage of slides within each lesion category that show significant composition differences relative to the healthy reference. Bars indicate the proportion of slides exceeding the Aitchison-distance significance thresholds for cell-type, structural, both, or one slide having significance with either composition. Error bars represent the uncertainty estimated by recalculating the percentages leaving out on slide at a time, showing how sensitive the results are to the inclusion of any individual slide.

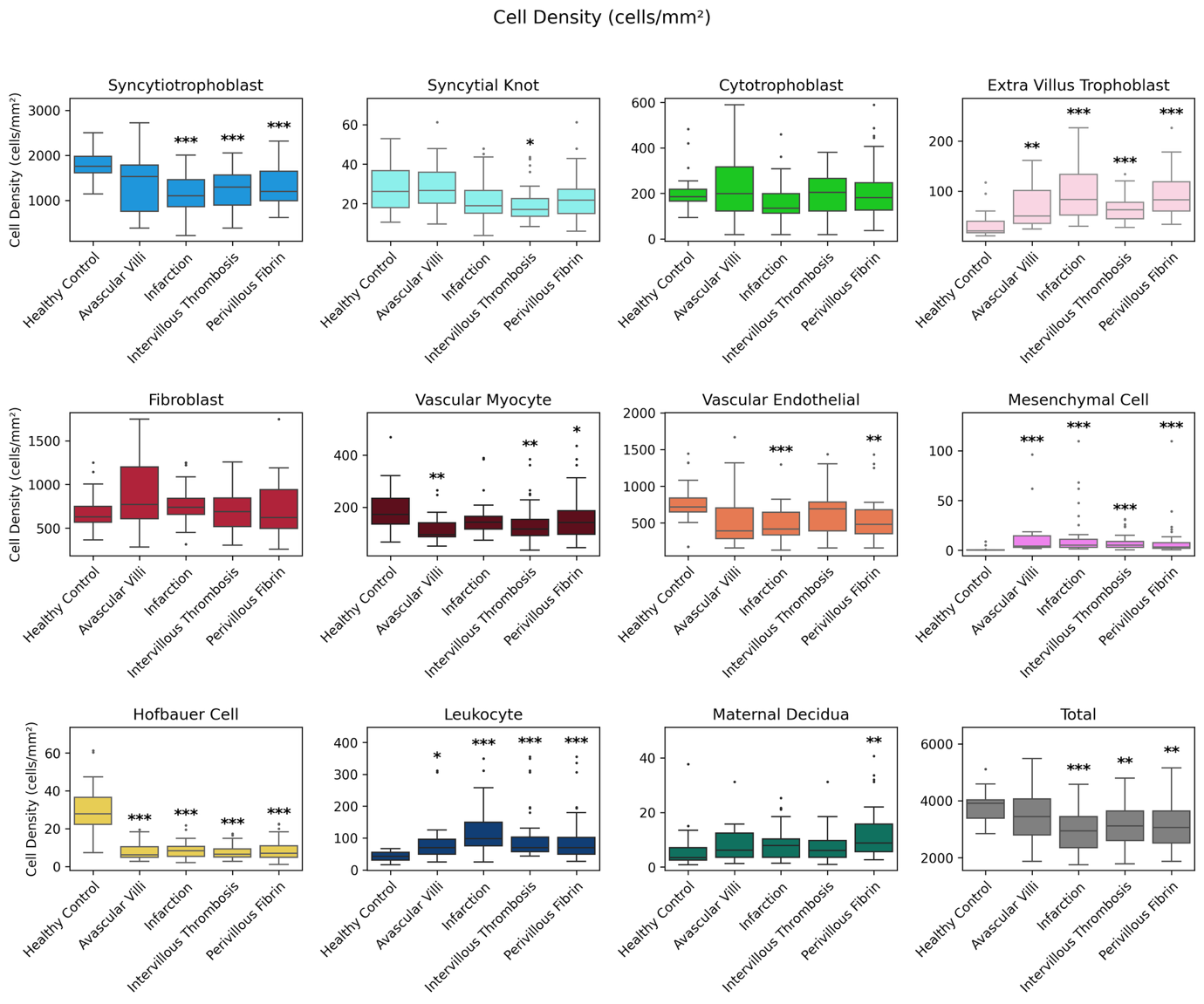

**Supplementary Figure 5:** Cell density (cells/mm²) across lesion categories for all cell types, including the total cell density across the slide. For each cell type, boxplots depict the median and interquartile range (IQR) of observed densities within each lesion group, with whiskers extending to 1.5IQR and points representing outliers. Slides are included in each lesion category if that slide contains the corresponding lesion type. Statistical differences of each lesion type compared to the healthy control group were assessed using the Mann-Whiteney U test with Bonferroni correction. Significance levels: ***** p<0.05, ****** p<0.01, ******* p<0.001.

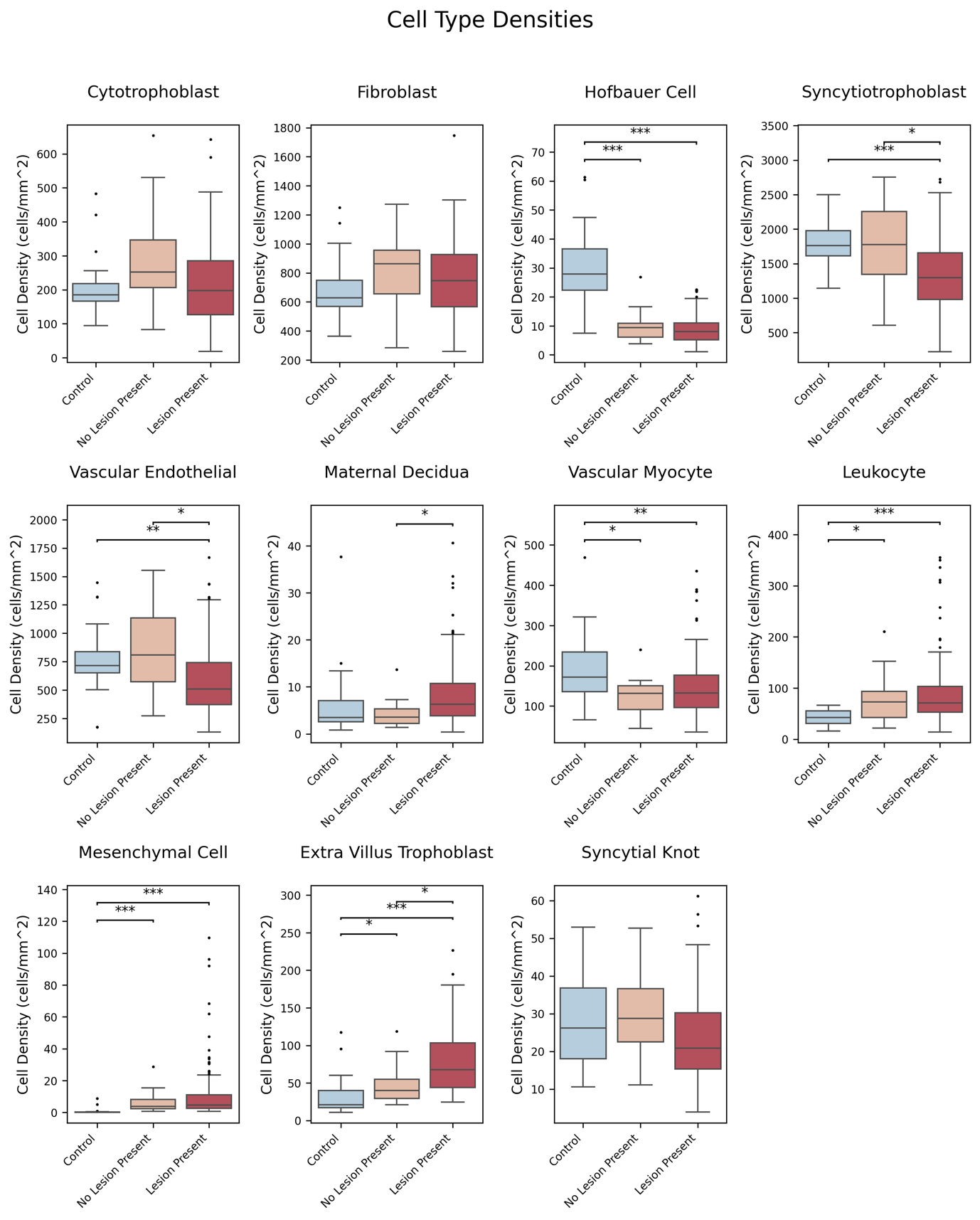
**Supplementary Figure 6:** Cell density (cells/mm²) across lesion categories for slide types: healthy control slides, slides from placentas with lesions but without a lesion on that sampled slide (“No Lesion Present”), and slides from those placentas on which a lesion is present (“Lesion Present”). For each cell type, boxplots depict the median and interquartile range (IQR) of observed densities within each lesion group, with whiskers extending to 1.5IQR and points representing outliers. Slides are included in each lesion category if that slide contains the corresponding lesion type. Statistical differences of each lesion type compared to the healthy control group were assessed using the Mann-Whiteney U test with Bonferroni correction. Significance levels: ***** p<0.05, ****** p<0.01, ******* p<0.001.

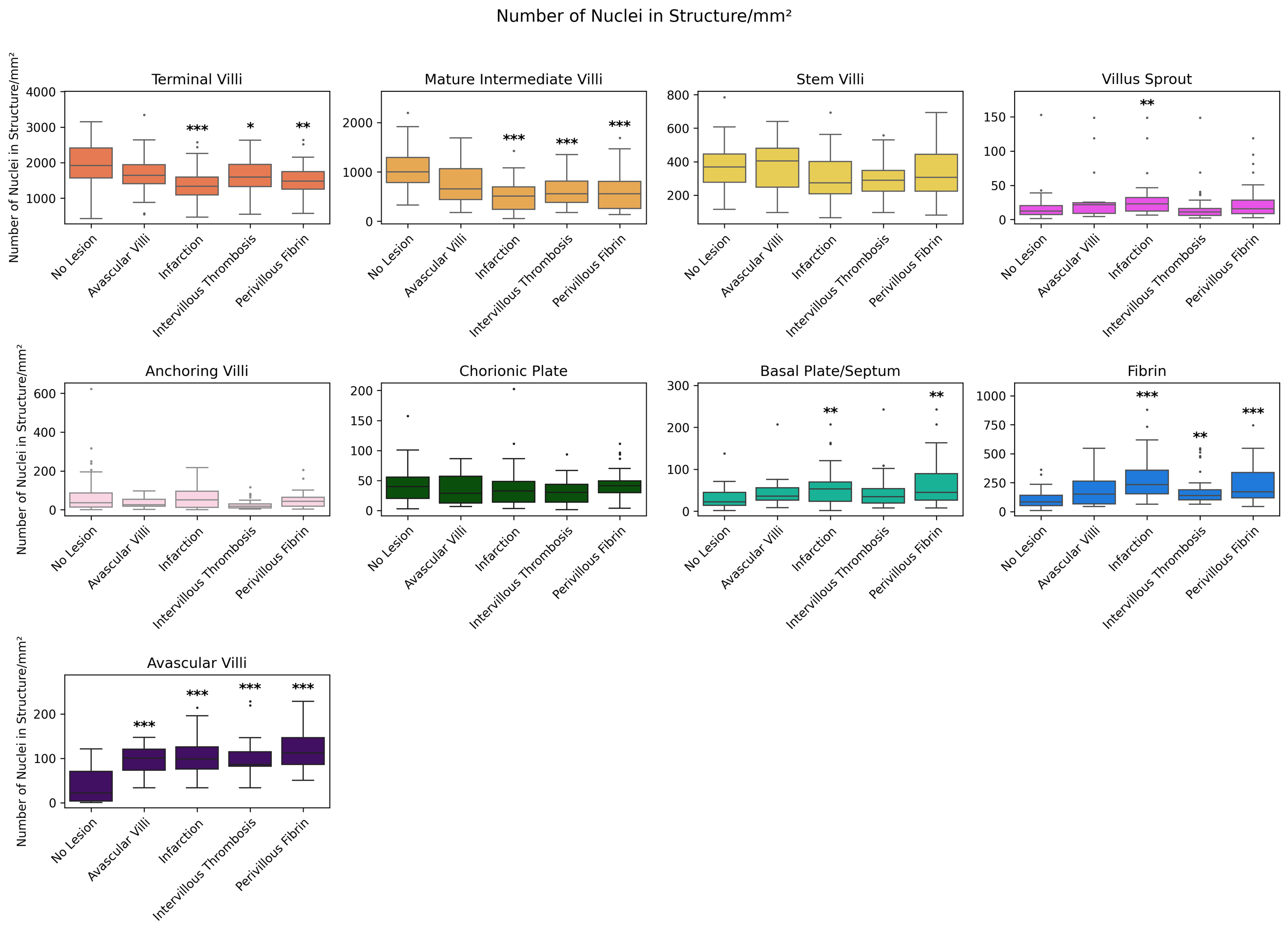

**Supplementary Figure 7:** Number of nuclei in each structure (cells/mm²) across lesion categories for all structure types. For each structure, boxplots depict the median and interquartile range (IQR) of observed densities within each lesion group, with whiskers extending to 1.5IQR and points representing outliers. Slides are included in each lesion category if that slide contains the corresponding lesion type. Statistical differences of each lesion type compared to the healthy control group were assessed using the Mann-Whiteney U test with Bonferroni correction. Significance levels: ***** p<0.05, ****** p<0.01, ******* p<0.001. Notably, basal plate densities differ significantly in slides with infarction and perivillous fibrin deposition, motivating the decision to focus subsequent analyses on the parenchyma region.

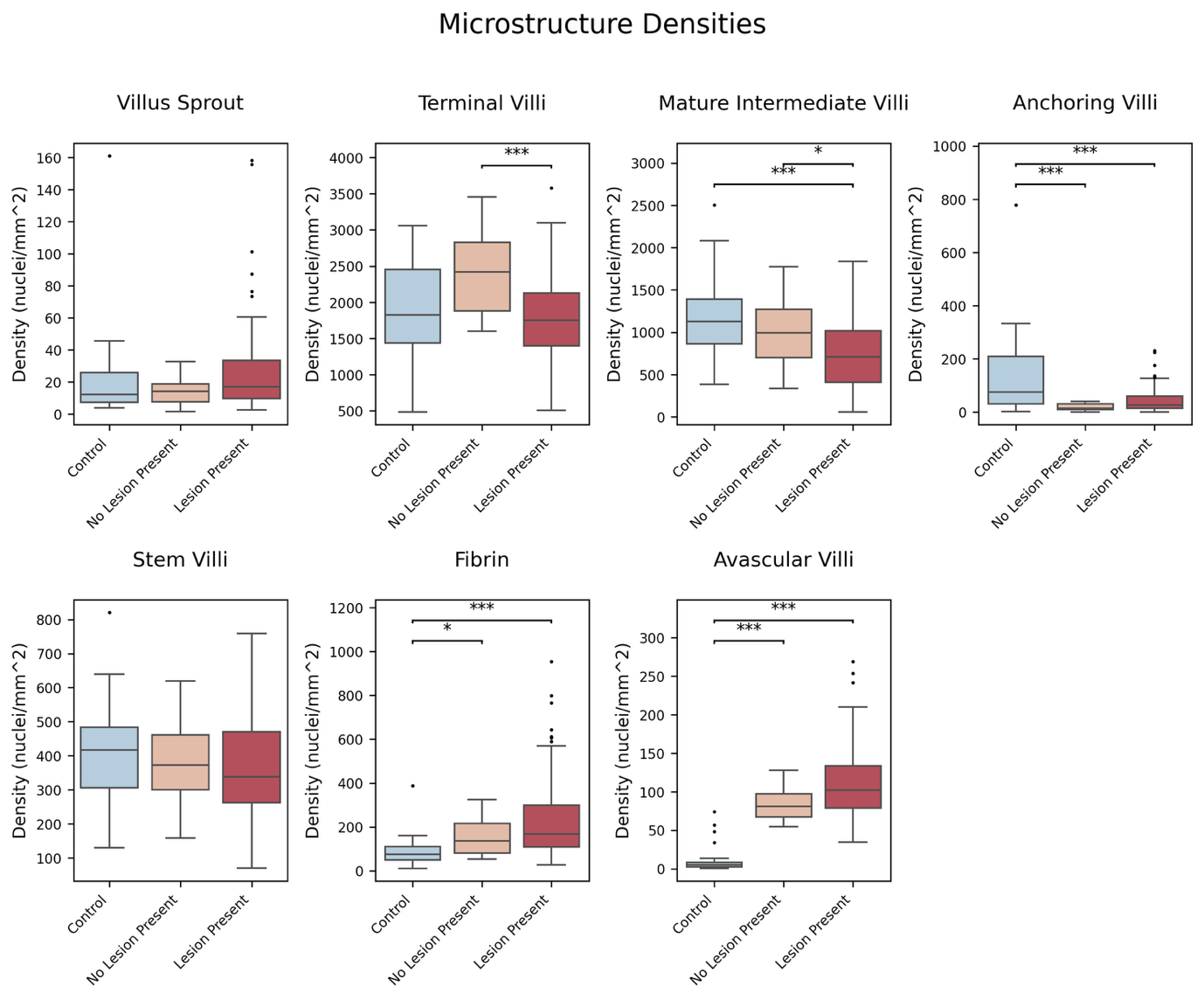

**Supplementary Figure 8:** Number of nuclei in each structure (cells/mm²) across lesion categories for for slide types: healthy control slides, slides from placentas with lesions but without a lesion on that sampled slide (“No Lesion Present”), and slides from those placentas on which a lesion is present (“Lesion Present”). For each structure, boxplots depict the median and interquartile range (IQR) of observed densities within each lesion group, with whiskers extending to 1.5IQR and points representing outliers. Statistical differences of each lesion type compared to the healthy control group were assessed using the Mann-Whiteney U test with Bonferroni correction. Significance levels: * p<0.05, ** p<0.01, *** p<0.001.

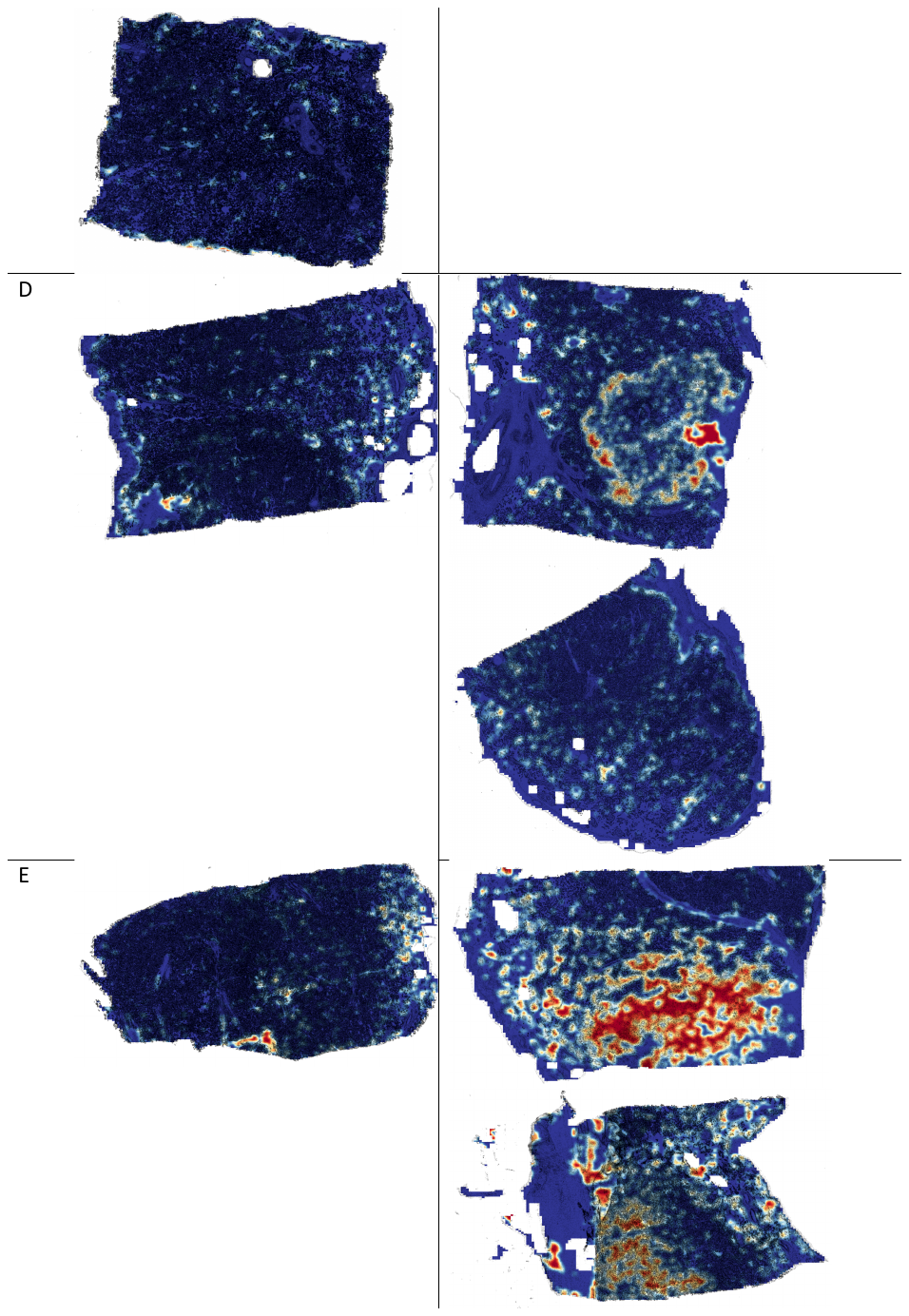

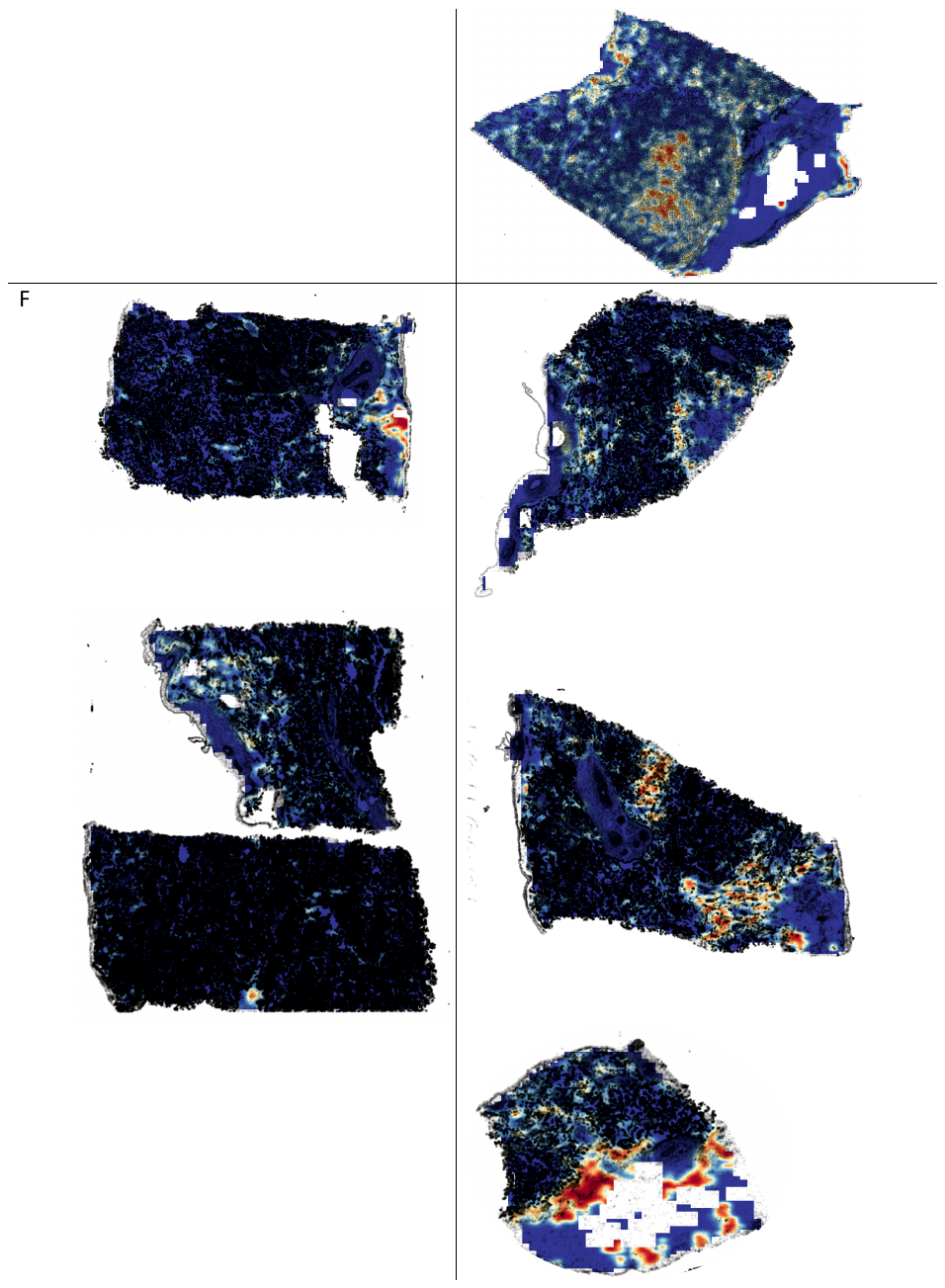

**Supplementary Figure 9:** Visualisation of the spatial distribution of fibrin using local autocorrelation maps for all placentas with infarction that also have paired “no apparent lesion” slides. Each panel shows local autocorrelation values, where higher values (red) indicate dense, spatially clustered fibrin and lower values (blue) reflect sparse or dispersed fibrin. Black dots mark the locations of individual nuclei. Comparing lesion-present and no-lesion slides demonstrates the pronounced increase in locally clustered fibrin deposition surrounding infarcted regions.

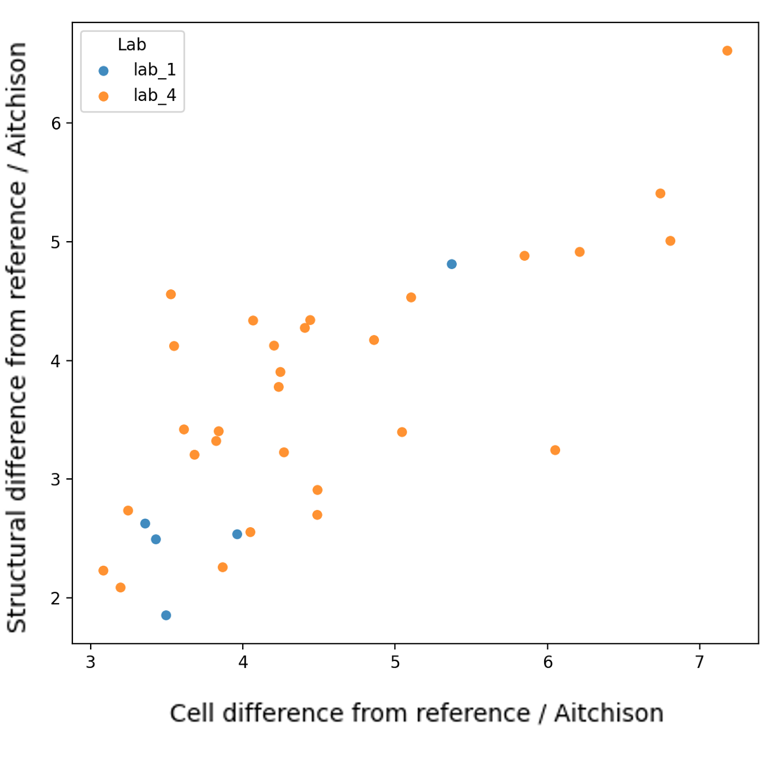

**Supplementary Figure 10:** Assessment of potential batch effects in slides with infarction. Each point represents an infarction-positive slide, showing its Aitchison distance from the healthy reference for cell-type composition and structural composition. Points are coloured by laboratory of origin. The overlap in distribution between labs and the absence of systematic separation along either axis suggest no strong batch effect between laboratories for infarction slides.

**Supplementary Information**

Supplementary Table 1

**Supplementary Table 1:** Summary of lesion types present across all slides. Single-lesion categories list the number of slides containing only that lesion type. Multiple lesion types category includes slides that contain more than one lesion; each specific combination of co-occurring lesions is shown in italics beneath this heading, with the corresponding number of slides for each combination.

| **Lesion Types Present** | **Number of Slides** |
| --- | --- |
| Perivillous Fibrin | 22 |
| Infarction | 18 |
| Intervillous Thrombosis | 18 |
| Avascular Villi | 6 |
| Multiple Lesion Types | 31 |
| *Intervillous Thrombosis + Perivillous Fibrin* | 10 |
| *Infarction + Perivillous Fibrin* | 7 |
| *Infarction + Intervillous Thrombosis* | 5 |
| *Avascular Villi + Intervillous Thrombosis + Perivillous Fibrin* | 3 |
| *Avascular Villi + Infarction* | 2 |
| *Avascular Villi + Perivillous Fibrin* | 2 |
| *Avascular Villi + Infarction + Perivillous Fibrin* | 1 |
| *Avascular Villi + Intervillous Thrombosis + Infarction* | 1 |

**Supplementary Table 2:** Cohen's d effect sizes quantifying differences in cell-type densities between placental lesion groups and healthy control placentas.

| **Cell Type** | **Avascular Villi** | **Infarction** | **Perivillous Fibrin** | **Intervillous Thrombosis** |
| --- | --- | --- | --- | --- |
| Syncytiotrophoblast | -0.89 | -1.58 | -1.22 | -1.41 |
| Syncytial Knot | 0.11 | -0.49 | -0.45 | -0.72 |
| Cytotrophoblast | 0.22 | -0.47 | 0.03 | -0.19 |
| Extra Villus Trophoblast | 1.00 | 1.42 | 1.55 | 1.21 |
| Fibroblast | 0.61 | 0.35 | 0.10 | 0.10 |
| Vascular Myocyte | -0.95 | -0.58 | -0.54 | -0.71 |
| Vascular Endothelial | -0.54 | -1.09 | -0.85 | -0.43 |
| Mesenchymal Cell | 0.94 | 0.75 | 0.55 | 1.15 |
| Hofbauer Cell | -1.93 | -2.33 | -2.38 | -2.55 |
| Leukocyte | 0.98 | 1.37 | 0.83 | 0.93 |
| Maternal Decidua | 0.30 | 0.34 | 0.64 | 0.16 |

**Supplementary Table 3:** Cohen's d effect sizes quantifying differences in structure-type densities between placental lesion groups and healthy control placentas.

| **Tissue Structure Type** | **Avascular Villi** | **Infarction** | **Perivillous Fibrin** | **Intervillous Thrombosis** |
| --- | --- | --- | --- | --- |
| Terminal Villi | -0.15 | -0.63 | -0.42 | -0.30 |
| Mature Intermediate Villi | -0.84 | -1.66 | -1.41 | -1.30 |
| Stem Villi | -0.10 | -0.45 | -0.30 | -0.55 |
| Villus Sprout | 0.40 | 0.32 | 0.15 | -0.07 |
| Anchoring Villi | -0.73 | -0.61 | -0.83 | -1.03 |
| Fibrin | 1.00 | 1.40 | 1.10 | 0.89 |
| Avascular Villi | 3.28 | 2.74 | 2.93 | 2.52 |

**Supplementary Table 4:** Cohen’s d effect sizes comparing cell-type densities abundances between diagnostic groups. Slides classified as no apparent lesion correspond to histopathologically normal placentas from patients with a recorded diagnosis, whereas lesion present indicates placentas with identifiable histopathological lesions.

| **Cell Type** | **No Apparent Lesion** | **Lesion Present** |
| --- | --- | --- |
| Syncytiotrophoblast | 0.0083 | -0.93 |
| Syncytial Knot | 0.22 | -0.36 |
| Cytotrophoblast | 0.71 | 0.11 |
| Extra Villus Trophoblast | 0.57 | 1.07 |
| Fibroblast | 0.64 | 0.32 |
| Vascular Myocyte | -0.99 | -0.69 |
| Vascular Endothelial | 0.32 | -0.61 |
| Mesenchymal Cell | 1.20 | 0.63 |
| Hofbauer Cell | -1.83 | -2.98 |
| Leukocyte | 1.10 | 0.81 |
| Maternal Decidua | -0.34 | 0.30 |

**Supplementary Table 5:** Cohen’s d effect sizes comparing structure-type densities abundances between diagnostic groups. Slides classified as no apparent lesion correspond to histopathologically normal placentas from patients with a recorded diagnosis, whereas lesion present indicates placentas with identifiable histopathological lesions.

| **Tissue Structure Type** | **No Apparent Lesion** | **Lesion Present** |
| --- | --- | --- |
| Terminal Villi | 0.78 | -0.21 |
| Mature Intermediate Villi | -0.38 | -1.19 |
| Stem Villi | -0.15 | -0.30 |
| Villus Sprout | -0.31 | 0.11 |
| Anchoring Villi | -0.92 | -1.22 |
| Fibrin | 0.81 | 0.84 |
| Avascular Villi | 3.56 | 2.31 |
